# Neurophysiological correlates of short-term recognition of sounds: Insights from magnetoencephalography

**DOI:** 10.1101/2023.12.07.570594

**Authors:** E. Serra, M. Lumaca, E. Brattico, P. Vuust, M. L. Kringelbach, L. Bonetti

**Affiliations:** Center for Music in the Brain, Department of Clinical Medicine, Aarhus University & The Royal Academy of Music, Aarhus, Aalborg, Denmark; Centre for Eudaimonia and Human Flourishing, Linacre College, University of Oxford, Oxford, United Kingdom; Department of Psychiatry, University of Oxford, Oxford, United Kingdom; Department of Education, Psychology, Communication, University of Bari Aldo Moro, Italy

**Keywords:** Short-term memory, Auditory memory, Music, Predictive coding (PC), Magnetoencephalography (MEG)

## Abstract

Understanding the brain’s dynamic retrieval and updating of encoded information is a key focus in memory research. This study employed a same versus different auditory paradigm to investigate short-term auditory recognition within a predictive coding (PC) framework, concerning the perceptual interplay between experience-informed predictions and incoming sensory information. Using magnetoencephalography (MEG), we captured the neurophysiological correlates associated with a single-sound, short-term memory task. Twenty-six healthy participants were tasked with recognizing whether presented sounds were same or different compared to strings of standard stimuli. To prompt conscious memory retention, a white noise interlude separated these sounds from the standards. MEG sensor-level results revealed that recognition of same sounds elicited two significantly stronger negative components of the event-related field compared to different sounds. The first one was the N1, peaking 100ms post-sound onset, while the second one corresponded to a slower negative component arising between 300 and 600ms after sound onset. This effect was observed in several significant clusters of MEG sensors, especially temporal and parietal regions of the scalp. Conversely, different sounds produced scattered and smaller clusters of stronger activity than same sounds, peaking later than 600ms after sound onset. Source reconstruction using beamforming algorithms revealed the involvement of auditory cortices, hippocampus, and cingulate gyrus in both conditions. Overall, the results are coherent with PC principles and previous results on the brain mechanisms underlying auditory recognition, highlighting the relevance of early and later negative brain responses for successful prediction of previously listened sounds in the context of conscious short-term memory.

## Introduction

Unravelling the intricate neural mechanisms underlying memory function in humans, namely how the brain encodes, processes, and recovers information from its dynamic environment, constitutes a central focus in cognitive neuroscience. Traditionally, various facets of memory have been explained through frameworks such as the multi-store model, which illustrates the movement of information from sensory to short-term memory, and onward to long-term memory through rehearsal (Atkinson & Shiffrin, 1977), and the working memory model, which delineates the active processing within short-term memory via the central executive, phonological loop, visuospatial sketchpad, and episodic buffer (Baddeley & Hitch, 1974; Berz, 1995; Caclin & Tillmann, 2018). More recently, it has been proposed that memory functions could be studied in the context of information processing theories (Vecchi & Gatti, 2020). Among such theories, predictive coding (PC) stands out as a well-established framework which explains how our brain efficiently interacts with sensory scenes (Friston, 2005). PC suggests that the brain acts as a generative system, comparing internally generated predictions shaped by templates of previously encoded stimuli against incoming sensory data. Prediction errors, arising from discrepancies between top-down predictions and bottom-up sensory inputs, drive the updating of expectations in the neural hierarchy. This continual integration of anticipation and processing forms a Bayesian-like heuristic framework in PC (Friston, 2018).

The study of predictive processes in sensory memory spans both automatic and conscious levels of perception. To investigate these processes, various iterations of the oddball paradigm and same/different tasks are frequently employed. In the oddball paradigm, the regular flow of recurring stimuli is intermittently interrupted by infrequent deviants, drawing attention to novel or unexpected elements in the sensory environment (Squires et al., 1976; Duncan-Johnson & Donchin, 1977). This paradigm provides a valuable tool for probing automatic processing in sensory memory, as the deviation from the expected pattern elicits distinct neural responses. Similarly, same/different tasks involve the comparison of stimuli to discern similarities or differences, offering insights into conscious engagement with sensory information (Näätänen, 2018; Ulanovsky, Las & Nelken, 2003). In this regard, studies utilizing PC paradigms in non-verbal auditory contexts (Bonetti et al., 2018; Lumaca et al., 2019b; Quiroga-Martinez et al., 2021, 2022) have offered a specialized avenue for dissecting such perceptual and cognitive components in automatic and conscious processing. Importantly, the domain of musical paradigms has emerged as a valuable terrain for understanding the interplay between auditory perception, cognitive abilities, and long-term training, exposing their neural underpinnings (Bonetti & Costa, 2016, 2018, 2019; Bonetti et al., 2021b; Costa, 2013; Costa et al., 2004; Crisculo et al., 2019, 2022; Husain et al., 2002; Iorio et al., 2022; Moreno et al., 2011, 2009; Pando-Naude et al., 2021; Schellenberg, 2004, 2006, 2011). In this interdisciplinary landscape, the application of PC paradigms to investigate non-linguistic auditory processing aligns with the broader exploration of predictive processes, and their potential interactions with cognitive functions, including memory.

Pairing these types of studies with neuroscientific techniques which possess fine-grained temporal resolution like EEG and MEG, exposes unique fast-scale neural responses occurring in response to standard and discrepant stimuli (Hillebrand et al., 2018). In fact, M/EEG measurements reveal an array of contiguous automatic event-related potentials/fields (ERP/Fs) categorized into three main classes based on their latencies (Assecondi, Villa-Sánchez & Shapiro, 2022). When contextualising their succession and distinct topographies in relation to PC, it becomes evident that auditory information processing provokes spatially distributed neural responses which cross neural hierarchies at distinct processing stages. Firstly, short-latency ERPs like P1, N1, mismatch negativity (MMN), and P2 express early sensory and perceptual processes (Fogarty, Barry & Steiner, 2020). N1 and MMN, within pre-attentive neural responses, garner particular attention in PC paradigms (Bonetti et al., 2022a). N1 peaks 100-150 ms after target onset across stimulus-contingent sensory areas as well as parietal (Molholm et al., 2006) and frontal regions (Giard et al., 1994), and is associated with initial feature extraction and discrimination of sensory inputs (Näätänen & Picton, 1987). Furthermore, N1 is known to decrease in amplitude with decreasing intensity of the stimulus, playing a crucial role in early sensory processing (Bruneau et al., 1993). MMN is expressed mainly in primary sensory areas such as the auditory cortex, and primarily reflects a change in the established regularity of auditory stimuli (Lumaca et al., 2019a; Liebenthal et al., 2003; Molholm et al., 2004; Opitz et al., 2002; Schönwiesner et al., 2007). As such, it is thought to index the activation of specialised automatic deviance detection networks (Kropotov et al., 1995; Bonetti et al., 2017, 2018, 2021c; 2022b, 2023; Lumaca et al. 2020). Studies consistently show heightened N1 amplitude in response to deviants, paralleling increased MMN in acoustic predictive processes (Näätänen, 2000; Inui et al., 2010). Oddball and same/different paradigm adaptations support the active cortical predictive process underlying these responses (Wacongne, Changeux & Dehaene, 2012; Heilbron & Chait, 2018; Caucheteux, Gramfort & King, 2023). On the other hand, studies of omission refute the notion that these responses result solely from dishabituation (Braga & Schönwiesner, 2022). Notably, N1 and MMN expression is modulated by task demands and target characteristics (Koelsch, Vuust & Friston, 2019). For instance, stimuli that receive active attention elicit a more robust response compared to their unattended counterparts (Lijffijt et al., 2009). Subsequently, mid-latency ERPs, like fronto-centrally and parietally expressed N2 and P3, reflect higher-order cognitive processes following initial perception (Polich, 2007; Rutiku et al., 2015; Assecondi, Villa-Sánchez & Shapiro, 2022). Finally, the fronto-parietally expressed negative slow wave (NSW), occurs between 500-1000 ms, and is associated with feedback mechanisms in PC, indicating top-down attentional allocation within memory recollection (Barascud et al., 2016; Fogarty, Barry & Steiner, 2020). Specifically, sustained negativity in this component is thought to signal top-down attentional allocation within a process of memory recollection (Mecklinger, 1998). Taken together, these distinct ERP classes elucidate the intricate neural processes associated with sensory prediction, attentional allocation, and memory recollection within the broader context of PC. As such, these stages can be broadly divided into early deviance detection responses which are primed by expectation, and subsequent cognitive processes linked to the updating of expectation through higher-order integration of prediction errors.

The complex nature of ERP manifestation emphasizes that understanding the roles of sound-induced ERPs within memory processes is a non-trivial task. Recently, we were able to explore, through detailed analysis of ERP topographical and temporal expression, the spatio-temporal dynamics underpinning long-term, global recognition of musical sequences. This revealed a large number of hierarchically organised brain regions which were responsible for generating predictions of the upcoming sounds of the melodies (Bonetti et al., 2020, 2021a, 2022a, 2022c; Fernandez-Rubio et al., 2022a, 2022b). These studies provided solid evidence of the neural mechanisms underlying predictive processes during long-term recognition of musical sounds. In contrast, the rapid spatiotemporal dynamics of recognition of sounds in the context of short-term, local encoding and retrieval remain in part elusive (Costa et al., 2023).

To this end, the present study introduces a novel same/different auditory recognition paradigm, incorporating a white noise element strategically placed between the encoding and recognition phases of same and different stimuli. This intentional addition serves a dual purpose: thwarting neuronal adaptation and promoting conscious engagement of short-term memory. Hence, the aims of the present study are twofold. Firstly, utilising MEG we aim to identify specific neurophysiological measures indicative of cognitive functioning associated with auditory information processing and expectation updating. Secondly, we seek to examine the alignment of these identified neural markers with the mechanisms proposed by the PC framework. If PC mechanisms are indeed at play in sensory perception and contingent upon short-term memory, we hypothesize that ERPs elicited by same and different stimuli will exhibit differential expression. These distinctions are anticipated to manifest as unique temporal and spatial patterns, reflecting perceptual inference grounded in memory traces and the integration of error detection. Through this multifaceted approach, we aim to shed light on the intricate interplay between neural processes, short-term memory, and the PC framework in the context of auditory cognition.

## Materials and methods

### Participants

The study included a total of 26 volunteers (M=13), with a mean age of 24.72 ± 4.79 years. The sample was recruited in Denmark and consisted of individuals from Western countries who reported normal hearing and had a homogenous educational background. The study was conducted in accordance with the ethical guidelines of the Declaration of Helsinki – Ethical Principles for Medical Research. Furthermore, it was approved by the Ethics Committee of the Central Denmark Region, De Videnskabsetiske Komitéer for Region Midtjylland (Ref 1-10-72-411-17). All participants provided informed consent before participating in the experimental procedures.

### Experimental stimuli and design

To investigate the neural correlates of conscious auditory short-term recognition of sounds, we used a same/different auditory paradigm, similar to the ones reported by Albouy and colleagues (2017) and Costa and colleagues (2023), during magnetoencephalography (MEG) recordings.

The same/different paradigm consisted of three phases: *encoding (i)*, *retention (ii)*, *recognition (iii)* (Figure 1b). During the *encoding phase (i)*, all participants listened to a sequence of three identical sounds with the same pitch. Each of the three sounds lasted approximately 250 ms and was separated by the other sounds by a silence lasting approximately 1500 ms. During the *retention phase (ii)*, a 2000 ms white noise stimulus was introduced. Participants were specifically instructed to hold in memory the sound they had previously encountered during the *encoding phase (i)*. This interference was deliberately employed to ensure that participants actively maintained the original auditory information in their mind, thereby engaging in an active short-term memory process and diverging from a purely automatic response. In the third and final *recognition phase (iii)*, participants were presented with an additional sound that could have the same pitch or a different pitch than the three sounds they had heard during the *encoding phase (i)*. Participants were required to indicate, pressing a button with the left or right hand, whether the fourth sound matched the sounds presented during the *encoding phase (i)* or differed from them. Half of the participants were instructed to press the right button for the same sounds and the left button for the different sounds, while the other half were asked to reverse this. The target sound for the neural data was the one presented in the *recognition phase (iii)*.

**Figure 1.**
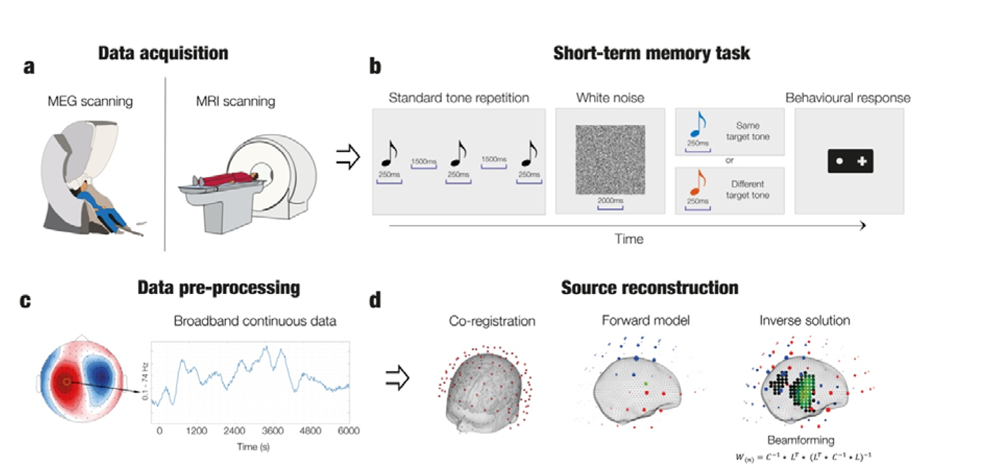
Experimental design, stimuli, and analysis pipeline. **a** – The brain activity of 26 volunteers was collected using magnetoencephalography (MEG) while they performed a same/different auditory recognition task. **b** – A tone was repeated three times for 250ms, separated by pauses of 1500ms. The third tone was followed by a 2000ms auditory white noise distractor, during which participants were asked to remember the tone. Subsequently, either the same or a different tone was played for 250ms, and participants were instructed to respond with a button press whether the tone was the same as the memorised tone or different. The order of presentation for same (n=20) and different (n=20) tasks was randomised. **c** – The continuous neural data was pre-processed to reduce external interference and eliminate internally generated artifacts. **d** – Source reconstruction analyses were conducted to identify brain sources which generated neural activity. First, MEG data was co-registered with individual or template anatomical MRI maps. Subsequently, MEG data was source reconstructed by computing an overlapping-spheres forward model using an 8-mm grid and calculating the inverse solution through a beamforming algorithm. Contrasts between neural responses to target same and different tones were calculated.

The volume of the stimuli was adjusted to a comfortable level for each participant (corresponding to +50 dB above the individual auditory threshold), and a few trial runs were conducted to ensure that participants understood the task. After this, the actual experiment, consisting of 40 trials, started.

### Data acquisition

The MEG data for this study was collected using a 306-channel Elekta Neuromag TRIUX MEG scanner (Elekta Neuromag, Helsinki, Finland) placed in a shielded room at Aarhus University Hospital in Denmark. The data was recorded at a sampling rate of 1000 Hz with a 0.1-330 Hz analogue filter. The head shape and position of four Head Position Indicator (HPI) coils were recorded using a 3D digitizer (Polhemus Fastrak, Colchester, VT, USA) in relation to three anatomical landmarks (nasion and left and right preauricular points). This information was used later to co-register the MEG data with the MRI scans. The HPI coils continuously monitored the head position during the MEG recordings to correct for head movement. Cardiac and eye movement data was also recorded using bipolar electrodes (electrocardiography [ECG] and electrooculography [EOG]). Later, this was used to remove artifacts from the neural data.

The MRI scans were conducted on a CE-approved 3T Siemens MRI-scanner at the same hospital. We recorded data which consisted of structural T1 (mp2rage) with a spatial resolution of 1.0 x 1.0 x 1.0 mm and the following sequence parameters: echo time = 2.61 ms, repetition time = 2300 ms, reconstructed matrix size = 256 x 256, echo spacing = 7.6 ms, bandwidth = 290 Hz/Px.

The MEG and MRI recordings were collected on separate days.

### MEG data pre-processing

The raw MEG data, consisting of 204 planar gradiometers and 102 magnetometers, was pre-processed using MaxFilter to reduce external interference (Taulu & Simola, 2006). This procedure consisted of applying signal space separation (SSS) to the data, down-sampling it from 1000 Hz to 250 Hz and compensating the head movement recorded by the cHPI coils (default step size: 10 ms). Here, the correlation limit between inner and outer subspaces that were used to reject overlapping intersecting inner/outer signals during SSS corresponded to 0.98.

After being converted into Statistical Parametric Mapping (SPM) format, the data was further pre-processed and analysed in MATLAB (MathWorks, Natick, MA, USA). This was done using a combination of codes from the Oxford Centre for Human Brain Activity (OHBA) Software Library (OSL) (Woolrich et al., 2011), a freely available software that is built upon Fieldtrip (Oostenveld et al., 2011), FSL (Woolrich et al., 2009), and SPM (Penny et al., 2011) toolboxes, and of in-house-built codes (LBPD, https://github.com/leonardob92/LBPD-1.0.git). The continuous MEG data was examined visually to identify and remove large artifacts using the OSLview tool, with less than 0.1% of the collected data being removed. Independent component analysis (ICA) was then used to eliminate interference from eyeblinks and heartbeats in the brain data (Mantini et al., 2011). The original signal was decomposed into independent components, and the components related to eyeblinks and heart activity (between 2 and 5 per participant) were isolated and discarded. The remaining components were then used to rebuild the signal. Finally, the neural signal was epoched in 40 trials (20 old and 20 new). Here, we applied the baseline correction by removing the mean signal recorded in the baseline (100 ms before the onset of the target sound of the recognition phase) from the post-stimulus brain signal. Each trial lasted 1600 ms (1500 ms plus 100 ms of baseline time).

### Massive univariate tests and Monte-Carlo simulations (MCS)

To assess the neural differences associated to the short-term recognition of same vs different sounds, several univariate *t*-tests were calculated by contrasting the neural data underlying same vs. different sounds and then corrected for multiple comparisons by employing MCS. In agreement with many other MEG and electroencephalography (EEG) task studies (Gross et al., 2013; Hari et al., 2018), averaging over trials was performed for each condition to obtain two mean trials, one for the old condition and one for the new condition. The sum-root square was then used to combine each pair of planar gradiometers. After this, a t-test was calculated for each MEG channel and time-point in the 0-1000 second range (number of time-points = 166), comparing the two experimental conditions. The resulting matrix was reshaped and binarized according to the p-values and t-values obtained from the t-tests (threshold of .05) described above. The resulting 3D matrix consisted of 0s where the t-tests were not significant and 1s where they were significant. To correct for multiple comparisons, MCS was used. Specifically, first the clusters of 1s were identified and their size stored. Then, the elements of the original binary 3D matrix were permuted 1000 times. For each time, the maximum cluster size of 1s was identified, and the distribution of the 1000 maximum cluster sizes was created. The original clusters were then considered significant if their size was greater than the 99.9% of the maximum cluster sizes of the permuted data. This MCS procedure was performed for the absolute value of the signal recorded by the magnetometers. The analysis was computed when old sounds produced stronger neural signal than new sounds, and vice versa.

### MEG source reconstruction

To estimate the neural sources of the signal recorded using MEG, we used the widely adopted set of algorithms related to the beamforming method (Huang, Mosher & Leahy, 1999; Huang et al., 2003; Luckhoo, Brookes & Woolrich, 2014). Here, we employed an implementation developed upon a combination of in-house-built codes and codes provided by OSL, SPM, and FieldTrip.

These algorithms consisted of three main steps: (i) co-registering individual MEG with MRI data, (ii) designing a forward model and (iii) computing the inverse solution.

At first, each individual T1-weighted MRI scan was aligned with the standard Montreal Neurological Institute (MNI) brain template using an affine transformation. It was then co-registered to the MEG sensor data using the Polhemus head shape information and the three fiducial points measured during the MEG session.

The forward model is a theoretical model that represents each brain source as an active dipole (brain voxel) and describes how the strength of each dipole would be reflected in the MEG sensor measurements. In this study, magnetometer channels and an 8-mm grid were used, resulting in 3559 dipoles within the whole brain. After co-registering the individual structural T1 data with the fiducial points, the forward model was computed using the Single Shell method, which is described in detail by Nolte (2003). Unfortunately, in 13 cases, it was not possible to collect the MRI data due to participants attending only one out of the two sessions. In those cases, we computed the source reconstruction using a template (MNI152-T1 with 8-mm spatial resolution).

Afterwards, the inverse solution was calculated using beamforming, a popular and effective method in the field. This procedure involves the use of weights that are sequentially applied to the source locations in order to isolate the contribution of each source to the activity recorded by the MEG channels at each time-point of the recorded brain data.

### Neural sources of the brain activity underlying processing of same and different sounds

After reconstructing the sources of the MEG signal for each time-point, we focused on the significant time-windows emerged from the contrasts computed at MEG sensor level. First, we averaged the reconstructed brain activity over all the time-points of each time-window, independently for each participant. This was done to show the sources responsible for generating the signals associated with both same and different sounds (an example is depicted in Figure 2). Second, we wished to assess whether we could detect any significant difference in MEG source space when contrasting the reconstructed brain activity of same vs. different sounds. To this aim, independently for the selected time-windows, we computed a t-test between same versus different sounds for each brain voxel. Afterwards, we used a 3D cluster-based MCS (MCS, α = .003, MCS *p-value* = .001) to correct for multiple comparisons. In this process, we first determined the sizes of clusters of significant neighbouring brain voxels. Next, we conducted 1000 permutations of the original data and calculated the sizes of clusters of significant neighbouring permuted brain voxels for each permutation. This generated a reference distribution of the largest cluster sizes observed in the permuted data. Finally, we deemed the original clusters to be significant if their sizes were larger than 99.99% of the clusters in the reference distribution. Further details on this MCS procedure can be read in our previous studies (Bonetti et al., 2020, 2021a, 2022a, 2022c; Fernandez-Rubio et al., 2022a, 2022b).

**Figure 2.**
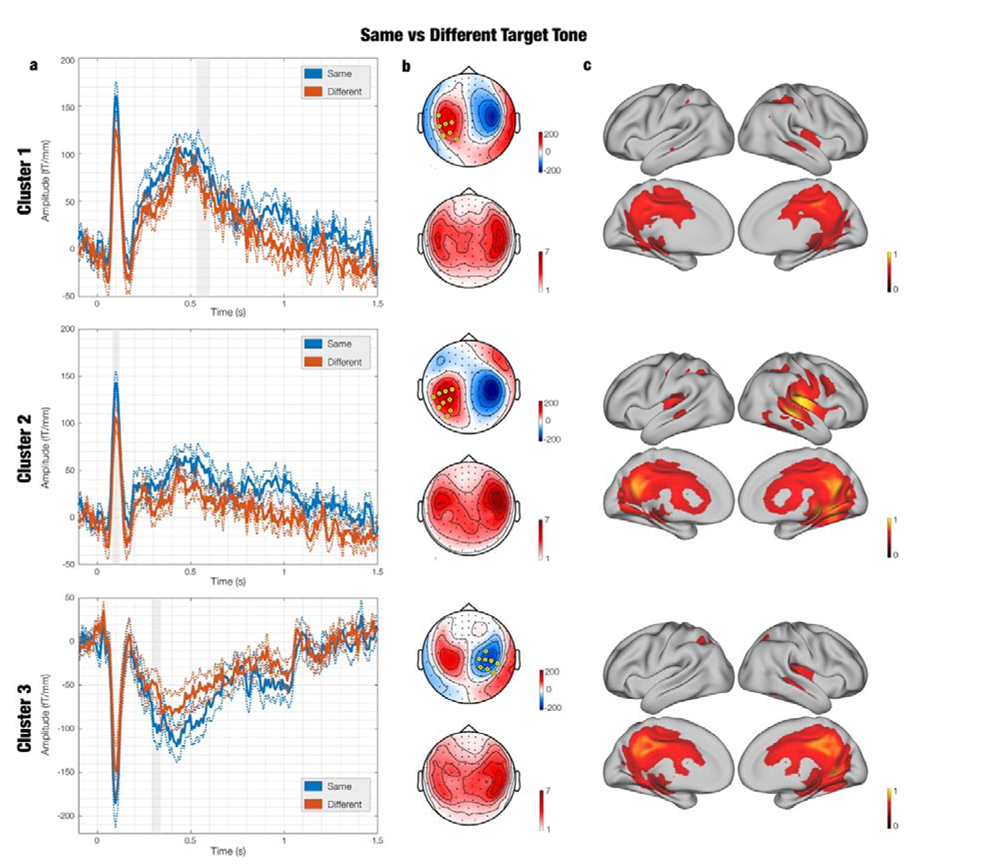
Neural responses to same versus different target tones: Temporal dynamics, scalp distribution, and cortical sources of significant clusters. **a** – MEG waveforms depicting the neural responses following onset of same (blue) and different (red) target tones in three most-significant clusters, averaged over participants. Dotted lines indicate the standard errors across participants. Gray shadows indicate the time-windows of significant differences between same and different timeseries **b** – Topographical maps showing the distribution of brain activity for gradiometers (top, fT/cm) and magnetometers (bottom, fT/cm) within the significant time-window emerged from MCS (MCS; α = .05, MCS p-value = .001). Both topoplots depict neural activity which was in same versus different target tone. Yellow circles within gradiometers show channel constituting significant clusters. **c** – Brain maps displaying the estimated neural sources of each for same versus different target tones. The values are t-statistics.

## Results

### Behavioural results

Before inspecting the neural data, we computed descriptive statistics of the behavioural performance at the task that participant completed during the MEG recording.

As expected, this analysis confirmed that participants could successfully perform the task. In fact, the same sounds were recognised on average 98.15% of the time, while the different sounds 95.41% of the time. In the same trials, reaction times were 969.50 ms ± 346.90 (mean ± standard deviation) for accurate responses and 1300.73 ms ± 356.00 for inaccurate responses. In the different trials, reaction times were 1025.01 ms ± 336.63 for accurate responses and 1121.33 ms ± 424.29 for inaccurate responses. No significant differences were observed for the accuracy and reaction times of the correctly identified same > different sounds (*p* > .025).

### Short-term recognition of single sounds

We contrasted the brain activity underlying same > different sounds using t-tests and MCS for multiple comparisons correction (t-test threshold = .05, MCS threshold = .001, 1000 permutations, see Methods for additional details). This analysis was conducted in the time-window (1000 ms) corresponding to the target sound in the *recognition phase (iii)* considered in the study (in Figure 2a we depicted the time-window starting from 100 ms before the onset of the sounds and ending 1500 ms after the onset, for illustrative purposes). This procedure returned a series of significant clusters. The strongest differences emerged for the contrast [same > different]. Key information on those clusters is reported in **Table 1**, while the main clusters are depicted in Figures 2b and **3b**. Additional details are provided in **Tables S1** and **S2.**

**Figure 3.**
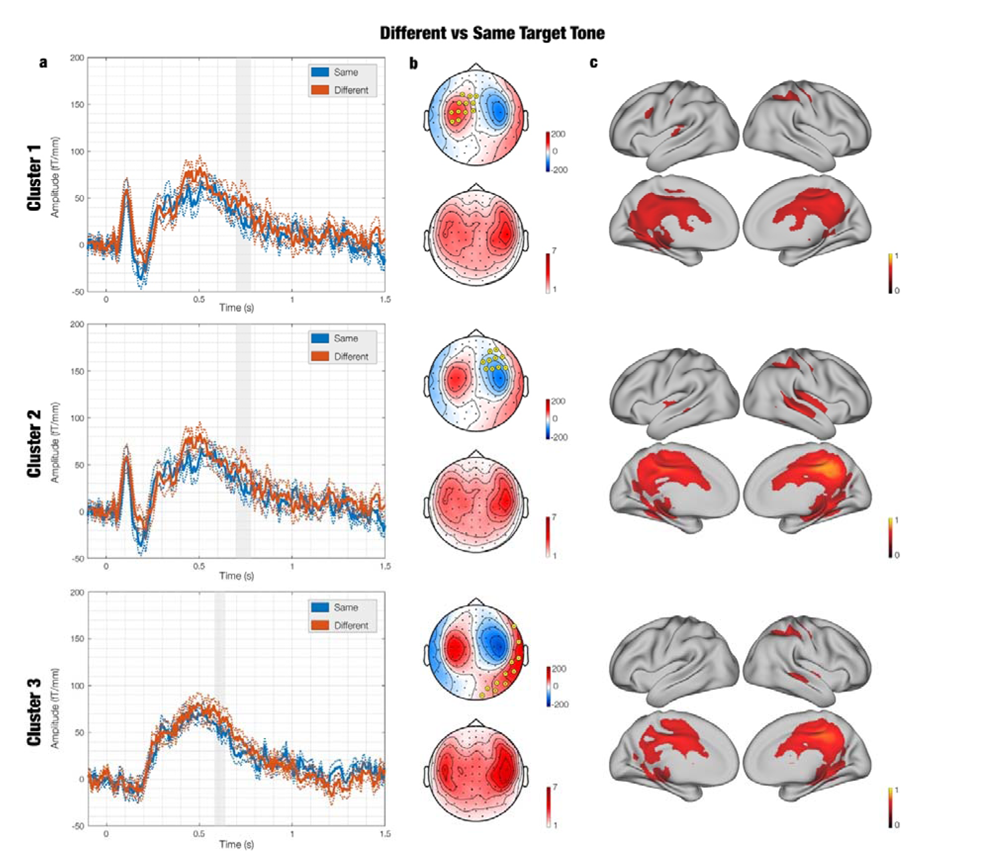
Neural responses to different versus same target tones: Temporal dynamics, scalp distribution, and cortical sources of significant clusters. **a** – MEG waveforms depicting the neural responses following onset of different (red) and same (blue) target tones in three most-significant clusters, averaged over participants. Dotted lines indicate the standard errors across participants. Gray shadows indicate the time-windows of significant differences between different and same timeseries **b** – Topographical maps showing the distribution of brain activity for gradiometers (top, fT/cm) and magnetometers (bottom, fT/cm) within the significant time-window emerged from MCS (MCS; α = .05, MCS p-value = .001). Both topoplots depict neural activity which was in different versus same target tone. Yellow circles within gradiometers show channel constituting significant clusters. **c** – Brain maps displaying the estimated neural sources of each for different versus same target tones. The values are t-statistics.

**Table 1.** Clusters of differential brain activity between same and different sounds (MEG sensors) Significant clusters obtained contrasting the neural activity at MEG sensor level underlying short-term recognition of same > different sounds. The table shows progressive cluster number, size (i.e., summation of the time-points and MEG channels involved), number of MEG channels forming the clusters, time-range for the significant difference, peak t-value within the cluster, MCS p-value.

| Cluster number | Size | MEG channels involved (n.) | Time-range (s) | Peak t-value | MCS p-value |
| --- | --- | --- | --- | --- | --- |
| *Same > Different* |  |  |  |  |  |
| 1 | 47 | 7 | 0.532 – 0.600 | 3.44 | < .001 |
| 2 | 46 | 9 | 0.084 – 0.116 | 4.14 | < .001 |
| 3 | 41 | 8 | 0.292 – 0.336 | 5.06 | < .001 |
| 4 | 31 | 9 | 0.768 – 0.800 | 4.53 | < .001 |
| 5 | 29 | 9 | 0.232 – 0.272 | 3.42 | < .001 |
| 6 | 28 | 5 | 0.884 – 0.920 | 3.73 | < .001 |
| 7 | 27 | 10 | 0.940 – 0.972 | 3.58 | < .001 |
| *Different > Same* |  |  |  |  |  |
| 1 | 91 | 12 | 0.704 – 0.780 | -4.49 | < .001 |
| 2 | 79 | 10 | 0.428 – 0.524 | -3.87 | < .001 |
| 3 | 68 | 11 | 0.584 – 0.648 | -4.67 | < .001 |
| 4 | 65 | 8 | 0.180 – 0.248 | -5.23 | < .001 |
| 5 | 53 | 8 | 0.620 – 0.688 | -4.62 | < .001 |
| 6 | 43 | 10 | 0.784 – 0.820 | -4.16 | < .001 |

### Neural sources of the MEG signal

The source reconstruction returned a network of brain areas that were active during short-term recognition of same and different sounds (Figures 2c and **3c**). This network comprised primary and secondary auditory cortices, brain regions of the hippocampal area and the posterior part of the medial cingulate gyrus. No significant differences emerged within same versus different and different versus same contrasts (*p* > .05).

## Discussion

The present study aimed to investigate the neural mechanisms underlying auditory short-term recognition in the context of the PC framework. We employed a same/different auditory paradigm comprising three key phases: *encoding*, *retention*, and *recognition*. In the *encoding* phase, participants listened to a sequence of three identical sounds with matching pitch. The subsequent *retention* phase introduced a 2000 ms white noise stimulus, prompting active retention of the initial auditory information. In the *recognition* phase, participants determined if a presented fourth sound matched or differed in pitch from those in the *encoding* phase. Our integration of this paradigm with MEG and MRI recordings allowed us to examine neurophysiological measures during the *recognition* phase, shedding light on cognitive processes related to auditory information processing, memory retrieval, and expectation updating in healthy adults.

On a behavioural level, there were no significant differences in response accuracy and reaction times between same and different target tones. This suggests that, irrespective of the neural modules engaged in their processing, there are no discernible temporal variations in the progression from sound perception to the selection of an appropriate response, whether it involves recognising same sounds or detecting different tones.

Interestingly, on a neural level, the same versus different contrast led to the emergence of significant clusters which primarily encompassed MEG sensors in temporal and parietal regions. It should be noted that this cluster category showed greater activity overall, suggesting that same target sounds elicited stronger neural responses compared to different target sounds. This finding is coherent with previous studies on the brain mechanisms underlying long-term (Bonetti et al., 2022, Fernández-Rubio et al., 2022) and short-term (Costa et al., 2023) recognition of auditory sequences. Consistently with those investigations, in the current study we reported an increased amplitude for the slow negative responses to the previously heard, same, sounds. However, such observed components peaked around 500 ms after sound onset, showing a latency which was considerably different from the 350-400 ms after each sound reported by Bonetti et al. (2022) and Fernández-Rubio et al. (2022). Moreover, in this experiment, we found a stronger amplitude of the N1 component for same sounds, a result that is usually not reported in these kinds of studies which typically see a stronger N1 activation in response to deviant sounds (Näätänen, 2000; Inui et al., 2010).

At the same time, our results also challenge some of the previous studies which reported attenuated neural signatures in response to predicted compared to unpredicted stimuli (Alink et al., 2010; den Ouden et al., 2010; Meyer, Travis & Olson, 2011). For instance, in our study the *same* condition exhibited a significantly enhanced N1 component compared to the different condition. This is in discordance with the typical observation in classical oddball paradigms, where N1 for standard stimuli is less pronounced than for deviants (May et al., 1999; Ranganath & Rainer, 2003; Grill-Spector, Henson & Martin, 2006). Such results have popularly been interpreted within the literature with respect to PC processes, leading to the proposition that N1 reduction in same conditions occurs as a result of facilitated auditory perception driven by expectation. The current study’s discrepant findings, however, should be considered in the context of its design and task demands, without automatic discreditation of PC mechanisms. In fact, given the explicit focus on conscious memorisation within the current design, and considering the breadth of factors which have shown to modulate short-latency responses, heightened expression of N1 may index feature extraction attuned to stimulus *relevance* rather than *deviance*. That is, a target tone which is the same as a successfully stored memory trace which must be recognised within incoming sensory information becomes more relevant for detection than deviant sounds. Here, we propose that increased N1 amplitude may reflect facilitated retrieval of task-relevant memory representations. This interpretation is further supported by the source-reconstructed topography of the N1 responses which originated not only in the auditory cortex, but also in the hippocampal area. Hippocampal activation indicates the involvement of memory structures in this process, as supported by similar studies outlining its role in music recognition (Burunat et al., 2014; Campo & Brattico, 2023). However, it is important to note that these target stimulus-dependent differences do not manifest at a behavioral level, likely due to a ceiling effect in the participants’ reaction times and accuracy, underscoring the need for future investigations.

On the other hand, the different versus same contrasts led to the emergence of smaller and scattered rather significant clusters of brain activity peaking after 600 ms from the onset of the target sounds. As for the recognition of the same sounds, this activity was observed in temporo-parietal regions of the scalp. Contrarily to the studies on long-term recognition of music which showed faster prediction error signals compared to responses indexing recognition and successful predictions, here we reported considerably slower activity showing enhanced prediction error responses (Bonetti et al., 2022b). This suggests that the violation of the representation of a single sound held in memory, as described in this paradigm, elicits a rather slow prediction error signal.

Finally, estimation of cortical generators of the components observed in each cluster uncovered an emergent network comprising primary and secondary auditory cortices, brain regions of the hippocampal area and the posterior part of the medial cingulate gyrus. These findings are in line with previous research, such as Kumar and colleagues’ (2016), who reported that the activity and connectivity between these three brain areas was crucial when holding sounds in mind. Nevertheless, in the present study, no significant difference in engagement of these areas can be reported between same and different conditions.

In conclusion, our study sheds light on the PC framework in the context of auditory short-term recognition by describing perceptual inference based on memory traces, and error detection integration. The findings provide novel interpretation for the neurophysiological mechanisms involved in auditory information processing of same sounds stored in memory traces, while elucidating prediction error computation and updating of expectation in response to different stimuli. Taken together, our findings suggest that negative components originated in the auditory cortices, medial temporal and parietal lobes are of primary importance for prediction and recognition processes of previously memorised sounds, both with regards to short-and long-term memory.

Nevertheless, it is important to note that the effect size of these results was not particularly strong. This could be attributed to methodological limitations rather than reflecting the true neurophysiological strength of expectation-based encoding and retrieval differences. Specifically, the conclusions drawn from the present study are constrained by the small sample size (n=26) and the use of standard neuroanatomical templates for the 13 participants who lacked MRI scans. This reliance on templates may have compromised the accuracy of source reconstruction, potentially explaining the absence of significant differences in neural source activation between the two stimulus categories, while the analyses on MEG sensor level provided solid and significant results.

Given the findings and limitations of the present study, it is crucial for future research to replicate these results using larger sample sizes. This would enhance the robustness and generatability of the findings. Furthermore, through subsequent investigations, this field will continue to mature the potential to provide invaluable insights into the underlying mechanisms governing predictive processes in conscious short-term recognition, and their intricate interplay with memory functions.

## Data availability

Please, find the codes used for the analyses reported in this study at the following link: https://github.com/leonardob92/MEG_STM_OneNote.git

Additional relevant in-house-built codes can be found at the following link: (LBPD, https://github.com/leonardob92/LBPD-1.0.git).

The multimodal neuroimaging data related to the experiment is available upon reasonable request.

## Acknowledgements

The Center for Music in the Brain (MIB) is funded by the Danish National Research Foundation (project number DNRF117).

ES is supported by the Luxembourg National Research Fund (FNR) (17906488) and the Society for Education, Music and Psychology Research (SEMPRE).

LB is supported by Carlsberg Foundation (CF20-0239), Lundbeck Foundation (Talent Prize 2022), Center for Music in the Brain, Linacre College of the University of Oxford (Lucy Halsall fund), and the Nordic Mensa Fund.

MLK is supported by Center for Music in the Brain and Centre for Eudaimonia and Human Flourishing, which is funded by the Pettit and Carlsberg Foundations.

We thank Simjon Radloff for his assistance during the data collection.

## Author contributions

LB and EB conceived the hypotheses. LB, EB and ES designed the study and the manuscript. LB, MLK, EB and PV recruited the resources for the experiment and writing up the manuscript. LB and EB collected the data. LB, ES and ML performed pre-processing and statistical analysis. MLK, PV, EB, ML and LB provided essential help to interpret and frame the results within the neuroscientific literature. ES and LB wrote the first draft of the manuscript. ES and LB prepared the figures. ML and EB edited the manuscript. All the authors contributed to and approved the final version of the manuscript.

## SUPPLEMENTARY TABLES

**Table S1.** Time-windows and channels of significant differences between same and different timeseries for three most significant clusters.

| Same vs Different |  |  |  |  |  |  |  |  |
| --- | --- | --- | --- | --- | --- | --- | --- | --- |
| Cluster 1 |  |  | Cluster 2 |  |  | Cluster 3 |  |  |
| MEG channel | Time 1 (start) | Time 2 (end) | MEG channel | Time 1 (start) | Time 2 (end) | MEG channel | Time 1 (start) | Time 2 (end) |
| MEG1631 | 0.53 | 0.60 | MEG1611 | 0.084 | 0.108 | MEG2221 | 0.29 | 0.33 |
| MEG1641 | 0.54 | 0.60 | MEG1621 | 0.084 | 0.108 | MEG2211 | 0.3 | 0.3 |
| MEG1621 | 0.54 | 0.60 | MEG1641 | 0.084 | 0.108 | MEG2441 | 0.3 | 0.33 |
| MEG1611 | 0.56 | 0.60 | MEG1631 | 0.084 | 0.108 | MEG2411 | 0.3 | 0.33 |
| MEG0241 | 0.58 | 0.60 | MEG1941 | 0.088 | 0.104 | MEG1131 | 0.3 | 0.3 |
| MEG1811 | 0.58 | 0.59 | MEG1521 | 0.092 | 0.116 | MEG2231 | 0.31 | 0.33 |
| MEG1911 | 0.60 | 0.60 | MEG1721 | 0.092 | 0.116 | MEG2431 | 0.33 | 0.33 |
|  |  |  | MEG1731 | 0.096 | 0.096 | MEG2421 | 0.33 | 0.34 |
|  |  |  | MEG1811 | 0.104 | 0.104 |  |  |  |

**Table S2.** Time-windows and channels of significant differences between different and same timeseries for three most significant clusters.

| Different vs Same |  |  |  |  |  |  |  |  |
| --- | --- | --- | --- | --- | --- | --- | --- | --- |
| Cluster 1 |  |  | Cluster 2 |  |  | Cluster 3 |  |  |
| MEG channel | Time 1 (start) | Time 2 (end) | MEG channel | Time 1 (start) | Time 2 (end) | MEG channel | Time 1 (start) | Time 2 (end) |
| MEG0421 | 0.704 | 0.78 | MEG1311 | 0.43 | 0.49 | MEG2131 | 0.584 | 0.616 |
| MEG0641 | 0.704 | 0.764 | MEG1121 | 0.43 | 0.52 | MEG2541 | 0.588 | 0.608 |
| MEG0411 | 0.716 | 0.764 | MEG1211 | 0.43 | 0.52 | MEG2331 | 0.592 | 0.616 |
| MEG0441 | 0.716 | 0.76 | MEG1231 | 0.43 | 0.52 | MEG2511 | 0.596 | 0.636 |
| MEG0331 | 0.716 | 0.764 | MEG1241 | 0.44 | 0.52 | MEG2521 | 0.596 | 0.644 |
| MEG0631 | 0.716 | 0.748 | MEG1031 | 0.44 | 0.44 | MEG1431 | 0.596 | 0.624 |
| MEG0231 | 0.728 | 0.744 | MEG1111 | 0.44 | 0.45 | MEG2621 | 0.596 | 0.64 |
| MEG0431 | 0.732 | 0.772 | MEG1321 | 0.45 | 0.46 | MEG2531 | 0.6 | 0.62 |
| MEG0711 | 0.736 | 0.744 | MEG1221 | 0.46 | 0.49 | MEG2641 | 0.6 | 0.648 |
| MEG0621 | 0.74 | 0.74 | MEG0931 | 0.52 | 0.52 | MEG2631 | 0.604 | 0.62 |
| MEG1621 | 0.748 | 0.748 |  |  |  | MEG1421 | 0.62 | 0.62 |
| MEG1811 | 0.748 | 0.748 |  |  |  |  |  |  |

## References

1. Alink, A., Schwiedrzik, C. M., Kohler, A., Singer, W., & Muckli, L. (2010). Stimulus predictability reduces responses in primary visual cortex. The Journal of Neuroscience, 30(8), 2960–2966. 10.1523/jneurosci.3730-10.2010

2. Albouy, P., Weiss, A., Baillet, S., & Zatorre, R. J. (2017). Selective entrainment of theta oscillations in the dorsal stream causally enhances auditory working memory performance. Neuron, 94(1), 193–206. e195. 10.1016/j.neuron.2017.03.015

3. Assecondi, S., Villa-Sánchez, B., & Shapiro, K. (2022). Event-related potentials as markers of efficacy for combined working memory training and transcranial direct current stimulation regimens: A proof-of-concept study. Frontiers in Systems Neuroscience, 16. 10.3389/fnsys.2022.837979

4. Atkinson, R. C., & Shiffrin, R. M. (1977). Human memory: A proposed system and its control processes. Human Memory, 7–113. 10.1016/b978-0-12-121050-2.50006-5

5. Baddeley, A. D., & Hitch, G. (1974). Working memory. Psychology of Learning and Motivation, 47–89. 10.1016/s0079-7421(08)60452-1

6. Barascud, N., Pearce, M. T., Griffiths, T. D., Friston, K. J., & Chait, M. (2016). Brain responses in humans reveal ideal observer-like sensitivity to complex acoustic patterns. Proceedings of the National Academy of Sciences, 113(5). 10.1073/pnas.1508523113

7. Berz, W. L. (1995). Working memory in music: A theoretical model. Music Perception, 12(3), 353–364. 10.2307/40286188

8. Bonetti, L., & Costa, M. (2016). Intelligence and musical mode preference. Empirical Studies of the Arts, 34(2), 160–176. 10.1177/0276237416628907

9. Bonetti, L., & Costa, M. (2018). Pitch-verticality and pitch-size cross-modal interactions. Psychology of Music, 46(3), 340–356. 10.1177/0305735617710734

10. Bonetti, L., & Costa, M. (2019). Musical mode and visual-spatial cross-modal associations in infants and adults. Musicae Scientiae, 23(1), 50–68. 10.1177/1029864917705001

11. Bonetti, L., Brattico, E., Bruzzone, S. E., Donati, G., Deco, G., Pantazis, D., Vuust, P., & Kringelbach, M. L. (2022a). Brain recognition of previously learned versus novel temporal sequences: A differential simultaneous processing. Cerebral Cortex, 33(9), 5524–5537. 10.1093/cercor/bhac439

12. Bonetti, L., Brattico, E., Carlomagno, F., Cabral, J., Stevner, A., Deco, G., Whybrow, P., Pearce, M., Pantazis, D., & Vuust, P. (2020). Spatiotemporal brain dynamics during recognition of the music of Johann Sebastian Bach. bioRxiv. 10.1101/2020.06.23.165191

13. Bonetti, L., Brattico, E., Carlomagno, F., Donati, G., Cabral, J., Haumann, N. T., Deco, G., Vuust, P., & Kringelbach, M. L. (2021a). Rapid encoding of musical tones discovered in whole-brain connectivity. NeuroImage, 245, 118735. 10.1016/j.neuroimage.2021.118735

14. Bonetti, L., Brattico, E., Vuust, P., Kliuchko, M., & Saarikallio, S. (2021b). Intelligence and music: lower intelligent quotient is associated with higher use of music for experiencing strong sensations. Empirical Studies of the Arts, 39(2), 194–215. 10.1177/0276237420951414

15. Bonetti, L., Bruzzone, S. E. P., Paunio, T., Kantojärvi, K., Kliuchko, M., Vuust, P., Palva, S., & Brattico, E. (2023). Moderate associations between BDNF VAL66MET gene polymorphism, musical expertise, and mismatch negativity. Heliyon, 9(5). 10.1016/j.heliyon 2023.e15600

16. Bonetti, L., Bruzzone, S. E. P., Sedghi, N. A., Haumann, N. T., Paunio, T., Kantojärvi, K., Kliuchko, M., Vuust, P., & Brattico, E. (2021c). Brain predictive coding processes are associated to COMT gene val158met polymorphism. NeuroImage, 233, 117954. 10.1016/j.neuroimage.2021.117954

17. Bonetti, L., Carlomagno, F., Kliuchko, M., Gold, B. P., Palva, S., Haumann, N. T., Tervaniemi, M., Huotilainen, M., Vuust, P., & Brattico, E. (2022b). Whole-brain computation of cognitive versus acoustic errors in music: A mismatch negativity study. Neuroimage: Reports, 2(4), 100145. 10.1016/j.ynirp.2022.100145

18. Bonetti, L., Fernández Rubio, G., Carlomagno, F., Pantazis, D., Vuust, P., & Kringelbach, M. L. (2022c). Revealing the spacetime hierarchical whole-brain dynamics of auditory predictive coding. bioRxiv, 2022-11. 10.1101/2022.11.19.517195

19. Bonetti, L., Haumann, N. T., Brattico, E., Kliuchko, M., Vuust, P., Särkämö, T., & Näätänen, R. (2018). Auditory sensory memory and working memory skills: Association between frontal MMN and performance scores. Brain Research, 1700, 86–98. 10.1016/j.brainres.2018.06.034

20. Bonetti, L., Haumann, N. T., Vuust, P., Kliuchko, M., & Brattico, E. (2017). Risk of depression enhances auditory pitch discrimination in the brain as indexed by the mismatch negativity. Clinical Neurophysiology, 128(10), 1923–1936. 10.1016/j.clinph.2017.07.004

21. Braga, A., & Schönwiesner, M. (2022). Neural substrates and models of omission responses and Predictive Processes. Frontiers in Neural Circuits, 16. 10.3389/fncir.2022.799581

22. Bruneau, N., Roux, S., Guérin, P., Garreau, B., & Lelord, G. (1993). Auditory stimulus intensity responses and frontal midline Theta Rhythm. Electroencephalography and Clinical Neurophysiology, 86(3), 213–216. 10.1016/0013-4694(93)90010-s

23. Burunat, I., Alluri, V., Toiviainen, P., Numminen, J., & Brattico, E. (2014). Dynamics of brain activity underlying working memory for music in a naturalistic condition. Cortex, 57, 254–269. 10.1016/j.cortex.2014.04.012

24. Caclin, A., & Tillmann, B. (2018). Musical and verbal short-term memory: Insights from Neurodevelopmental and neurological disorders. Annals of the New York Academy of Sciences, 1423(1), 155–165. 10.1111/nyas.13733

25. Campo, F. F., & Brattico, E. (2023). Remembering sounds in the brain: From locationist findings to Dynamic Connectivity Research. Rivista di Psicologia Clinica, (2), 7–39. 10.3280/rpc2-2022oa14002

26. Caucheteux, C., Gramfort, A., & King, J.-R. (2023). Evidence of a predictive coding hierarchy in the human brain listening to speech. Nature Human Behaviour, 7(3), 430–441. 10.1038/s41562-022-01516-2

27. Costa, M., Fine, P., & Ricci Bitti, P. E. (2004). Interval distributions, mode, and tonal strength of melodies as predictors of perceived emotion. Music Perception, 22(1), 1–14. 10.1525/mp.2004.22.1.1

28. Costa, M., Vuust, P., Kringelbach, M. L., & Bonetti, L. (2023). Age-related brain mechanisms underlying short-term recognition of musical sequences: An EEG study. bioRxiv. 10.1101/2023.03.12.532256

29. Criscuolo, A., Bonetti, L., Särkämö, T., Kliuchko, M., & Brattico, E. (2019). On the association between musical training, intelligence and executive functions in adulthood. Frontiers in psychology, 10, 1704. 10.3389/fpsyg.2019.01704

30. Criscuolo, A., Pando-Naude, V., Bonetti, L., Vuust, P., & Brattico, E. (2022). An ALE meta-analytic review of musical expertise. Sci Rep, 12(1), 11726. 10.1038/s41598-022-14959-4

31. den Ouden, H. E., Daunizeau, J., Roiser, J., Friston, K. J., & Stephan, K. E. (2010). Striatal prediction error modulates cortical coupling. The Journal of Neuroscience, 30(9), 3210–3219. 10.1523/jneurosci.4458-09.2010

32. Duncan-Johnson, C. C., & Donchin, E. (1977). On quantifying surprise: The variation of event-related potentials with subjective probability. Psychophysiology, 14(5), 456–467. 10.1111/j.1469-8986.1977.tb01312.x

33. Fernandez-Rubio, G., Brattico, E., Kotz, S. A., Kringelbach, M. L., Vuust, P., & Bonetti, L. (2022a). Magnetoencephalography recordings reveal the spatiotemporal dynamics of recognition memory for complex versus simple auditory sequences. Commun Biol, 5(1), 1272. 10.1038/s42003-022-04217-8

34. Fernández-Rubio, G., Carlomagno, F., Vuust, P., Kringelbach, M. L., & Bonetti, L. (2022b). Associations between abstract working memory abilities and brain activity underlying long-term recognition of auditory sequences. PNAS Nexus, 1(4), pgac216. 10.1093/pnasnexus/pgac216

35. Fogarty, J. S., Barry, R. J., & Steiner, G. Z. (2018). Sequential processing in the classic oddball task: ERP components, probability, and behavior. Psychophysiology, 56(3). 10.1111/psyp.13300

36. Fogarty, J. S., Barry, R. J., & Steiner, G. Z. (2020). Auditory stimulusL and responseLlocked ERP components and behavior. Psychophysiology, 57(5). 10.1111/psyp.13538

37. Friston, K. (2005). A theory of cortical responses. Philosophical Transactions of the Royal Society B: Biological Sciences, 360(1456), 815–836. 10.1098/rstb.2005.1622

38. Friston, K. (2018). Does predictive coding have a future? Nature Neuroscience, 21(8), 1019– 1021. 10.1038/s41593-018-0200-7

39. Giard, M. H., Perrin, F., Echallier, J. F., Thévenet, M., Froment, J. C., & Pernier, J. (1994). Dissociation of temporal and frontal components in the human auditory N1 wave: A scalp current density and dipole model analysis. Electroencephalography and Clinical Neurophysiology/Evoked Potentials Section, 92(3), 238–252. 10.1016/0168-5597(94)90067-1

40. Grill-Spector, K., Henson, R., & Martin, A. (2006). Repetition and the brain: Neural models of stimulus-specific effects. Trends in Cognitive Sciences, 10(1), 14–23. 10.1016/j.tics.2005.11.006

41. Gross, J., Baillet, S., Barnes, G. R., Henson, R. N., Hillebrand, A., Jensen, O., Jerbi, K., Litvak, V., Maess, B., Oostenveld, R., Parkkonen, L., Taylor, J. R., van Wassenhove, V., Wibral, M., & Schoffelen, J.-M. (2013). Good practice for conducting and Reporting Meg Research. NeuroImage, 65, 349–363. 10.1016/j.neuroimage.2012.10.001

42. Hari, R., Baillet, S., Barnes, G., Burgess, R., Forss, N., Gross, J., Hämäläinen, M., Jensen, O., Kakigi, R., Mauguière, F., Nakasato, N., Puce, A., Romani, G.-L., Schnitzler, A., & Taulu, S. (2018). IFCN-endorsed practical guidelines for clinical magnetoencephalography (MEG). Clinical Neurophysiology, 129(8), 1720–1747. 10.1016/j.clinph.2018.03.042

43. Heilbron, M., & Chait, M. (2018). Great expectations: Is there evidence for predictive coding in auditory cortex? Neuroscience, 389, 54–73. 10.1016/j.neuroscience.2017.07.061

44. Hillebrand, A., Gaetz, W., Furlong, P. L., Gouw, A. A., & Stam, C. J. (2018). Practical guidelines for clinical magnetoencephalography – another step towards best practice. Clinical Neurophysiology, 129(8), 1709–1711. 10.1016/j.clinph.2018.05.007

45. Huang, M. X., Mosher, J. C., & Leahy, R. M. (1999). A sensor-weighted overlapping-sphere head model and exhaustive head model comparison for Meg. Physics in Medicine and Biology, 44(2), 423–440. 10.1088/0031-9155/44/2/010

46. Huang, M. X., Shih, J. J., Lee, R. R., Harrington, D. L., Thoma, R. J., Weisend, M. P., Hanlon, F., Paulson, K. M., Li, T., Martin, K., Miller, G. A., & Canive, J. M. (2003). Commonalities and differences among vectorized beamformers in Electromagnetic Source Imaging. Brain Topography, 16(3), 139–158. 10.1023/b:brat.0000019183.92439.51

48. Husain, G., Thompson, W. F., & Schellenberg, E. G. (2002). Effects of musical tempo and mode on arousal, mood, and spatial abilities. Music Perception, 20(2), 151–171. 10.1525/mp.2002.20.2.151

49. Inui, K., Urakawa, T., Yamashiro, K., Otsuru, N., Takeshima, Y., Nishihara, M., Motomura, E., Kida, T., & Kakigi, R. (2010). Echoic memory of a single pure tone indexed by change-related brain activity. BMC Neuroscience, 11(1). 10.1186/1471-2202-11-135

50. Iorio, C., Brattico, E., Munk Larsen, F., Vuust, P., & Bonetti, L. (2022). The effect of mental practice on music memorization. Psychology of Music, 50(1), 230–244. 10.1177/0305735621995234

51. Koelsch, S., Vuust, P., & Friston, K. (2019). Predictive Processes and the Peculiar Case of Music. Trends in cognitive sciences, 23(1), 63–77. 10.1016/j.tics.2018.10.006

52. Kropotov, J. D., Nääänen, R., Sevostianov, A. V., Alho, K., Reinikainen, K., & Kropotova, O. V. (1995). Mismatch negativity to auditory stimulus change recorded directly from the Human Temporal Cortex. Psychophysiology, 32(4), 418–422. 10.1111/j.1469-8986.1995.tb01226.x

53. Kumar, S., Joseph, S., Gander, P. E., Barascud, N., Halpern, A. R., & Griffiths, T. D. (2016). A brain system for auditory working memory. The Journal of Neuroscience, 36(16), 4492–4505. 10.1523/jneurosci.4341-14.2016

54. Liebenthal, E., Ellingson, M. L., Spanaki, M. V., Prieto, T. E., Ropella, K. M., & Binder, J. R. (2003). Simultaneous ERP and fmri of the auditory cortex in a passive oddball paradigm. NeuroImage, 19(4), 1395–1404. 10.1016/s1053-8119(03)00228-3

55. Lijffijt, M., Lane, S. D., Meier, S. L., Boutros, N. N., Burroughs, S., Steinberg, J. L., Gerard Moeller, F., & Swann, A. C. (2009). P50, N100, and P200 sensory gating: Relationships with behavioral inhibition, attention, and working memory. Psychophysiology, 46(5), 1059–1068. 10.1111/j.1469-8986.2009.00845.x

56. Luckhoo, H. T., Brookes, M. J., & Woolrich, M. W. (2014). Multi-session statistics on beamformed Meg Data. NeuroImage, 95, 330–335. 10.1016/j.neuroimage.2013.12.026

57. Lumaca, M., Dietz, M. J., Hansen, N. Chr., QuirogaLMartinez, D. R., & Vuust, P. (2020). Perceptual learning of tone patterns changes the effective connectivity between Heschl’s gyrus and Planum Temporale. Human Brain Mapping, 42(4), 941–952. 10.1002/hbm.25269

58. Lumaca, M., Kleber, B., Brattico, E., Vuust, P., & Baggio, G. (2019a). Functional connectivity in human auditory networks and the origins of variation in the transmission of Musical Systems. eLife, 8. 10.7554/elife.48710

59. Lumaca, M., Trusbak Haumann, N., Brattico, E., Grube, M., & Vuust, P. (2019b). Weighting of neural prediction error by rhythmic complexity: A predictive coding account using mismatch negativity. European Journal of Neuroscience, 49(12), 1597–1609. 10.1111/ejn.14329

60. Mantini, D., Penna, S. D., Marzetti, L., de Pasquale, F., Pizzella, V., Corbetta, M., & Romani, G. L. (2011). A signal-processing pipeline for magnetoencephalography resting-state networks. Brain Connectivity, 1(1), 49–59. 10.1089/brain.2011.0001

61. May, P., Tiitinen, H., Ilmoniemi, R. J., Nyman, G., Taylor, J. G., & Näätänen, R. (1999). Frequency Change Detection in Human Auditory Cortex. Journal of Computational Neuroscience, 6(2), 99–120. 10.1023/a:1008896417606

62. Meyer, Travis, & Olson, C. R. (2011). Statistical learning of visual transitions in monkey inferotemporal cortex. Proceedings of the National Academy of Sciences, 108(48), 19401–19406. 10.1073/pnas.1112895108

63. Molholm, S., Martinez, A., Ritter, W., Javitt, D. C., & Foxe, J. J. (2004). The neural circuitry of pre-attentive auditory change-detection: An fmri study of pitch and duration mismatch negativity generators. Cerebral Cortex, 15(5), 545–551. 10.1093/cercor/bhh155

64. Molholm, S., Sehatpour, P., Mehta, A. D., Shpaner, M., Gomez-Ramirez, M., Ortigue, S., Dyke, J. P., Schwartz, T. H., & Foxe, J. J. (2006). Audio-visual multisensory integration in superior parietal lobule revealed by human intracranial recordings. Journal of Neurophysiology, 96(2), 721–729. 10.1152/jn.00285.2006

65. Moreno, S., Bialystok, E., Barac, R., Schellenberg, E. G., Cepeda, N. J., & Chau, T. (2011). Short-term music training enhances verbal intelligence and executive function. Psychol Sci, 22(11), 1425–1433. 10.1177/0956797611416999

66. Moreno, S., Marques, C., Santos, A., Santos, M., Castro, S. L., & Besson, M. (2009). Musical training influences linguistic abilities in 8-year-old children: more evidence for brain plasticity. Cereb Cortex, 19(3), 712–723. 10.1093/cercor/bhn120

67. Nolte, G. (2003). The magnetic lead field theorem in the quasi-static approximation and its use for magnetoencephalography forward calculation in realistic volume conductors. Physics in Medicine and Biology, 48(22), 3637–3652. 10.1088/0031-9155/48/22/002

68. Näätänen, R. (2018). Attention and brain function. Routledge.

69. Näätänen, R. (2000). Mismatch negativity (MMN): Perspectives for Application. International Journal of Psychophysiology, 37(1), 3–10. 10.1016/s0167-8760(00)00091-x

70. Näätänen, R., & Picton, T. (1987). The N1 wave of the human electric and magnetic response to sound: A review and an analysis of the component structure. Psychophysiology, 24(4), 375–425. 10.1111/j.1469-8986.1987.tb00311.x

71. Oostenveld, R., Fries, P., Maris, E., & Schoffelen, J.-M. (2011). FieldTrip: Open source software for advanced analysis of MEG, EEG, and invasive electrophysiological data. Computational Intelligence and Neuroscience, 2011, 1–9. 10.1155/2011/156869

72. Opitz, B., Rinne, T., Mecklinger, A., von Cramon, D. Y., & Schröger, E. (2002). Differential contribution of frontal and temporal cortices to auditory change detection: Fmri and ERP results. NeuroImage, 15(1), 167–174. 10.1006/nimg.2001.0970

73. Pando-Naude, V., Patyczek, A., Bonetti, L., & Vuust, P. (2021). An ALE meta-analytic review of top-down and bottom-up processing of music in the brain. Sci Rep, 11(1), 20813. 10.1038/s41598-021-00139-3

74. Penny, W. D., Friston, K. J., Ashburner, J. T., Kiebel, S. J., & Nichols, T. E. (2011). Statistical parametric mapping: The analysis of Functional Brain Images. Academic Press.

75. Polich, J. (2007). Updating p300: An integrative theory of P3A and p3b. Clinical Neurophysiology, 118(10), 2128–2148. 10.1016/j.clinph.2007.04.019

76. Ranganath, C., & Rainer, G. (2003). Neural mechanisms for detecting and remembering novel events. Nature Reviews Neuroscience, 4(3), 193–202. 10.1038/nrn1052

77. Rutiku, R., Martin, M., Bachmann, T., & Aru, J. (2015). Does the p300 reflect conscious perception or its consequences? Neuroscience, 298, 180–189. 10.1016/j.neuroscience.2015.04.029

78. Quiroga-Martinez, D. R., Hansen, N. C., Højlund, A., Pearce, M., Brattico, E., Holmes, E., Friston, K., & Vuust, P. (2021). Musicianship and melodic predictability enhance neural gain in auditory cortex during pitch deviance detection. Human Brain Mapping. 10.1101/2021.02.11.430838

79. QuirogaLMartinez, D. R., Basiński, K., Nasielski, J., Tillmann, B., Brattico, E., Cholvy, F., Fornoni, L., Vuust, P., & Caclin, A. (2022). Enhanced mismatch negativity in harmonic compared with Inharmonic sounds. European Journal of Neuroscience, 56(5), 4583– 4599. 10.1111/ejn.15769

80. Schellenberg, E. G. (2004). Music lessons enhance IQ. Psychol Sci, 15(8), 511–514. 10.1111/j.0956-7976.2004.00711.x

81. Schellenberg, E. G. (2006). Long-term positive associations between music lessons and IQ. Journal of educational psychology, 98(2), 457. 10.1037/0022-0663.98.2.457

82. Schellenberg, E. G. (2011). Music lessons, emotional intelligence, and IQ. Music Perception, 29(2), 185–194. 10.1525/mp.2011.29.2.185

83. Schönwiesner, M., Novitski, N., Pakarinen, S., Carlson, S., Tervaniemi, M., & Näätänen, R. (2007). Heschl’s gyrus, posterior superior temporal gyrus, and mid-ventrolateral prefrontal cortex have different roles in the detection of acoustic changes. Journal of Neurophysiology, 97(3), 2075–2082. 10.1152/jn.01083.2006

84. Squires, K. C., Wickens, C., Squires, N. K., & Donchin, E. (1976). The effect of stimulus sequence on the waveform of the cortical event-related potential. Science, 193(4258), 1142–1146. 10.1126/science.959831

85. Taulu, S., & Simola, J. (2006). Spatiotemporal signal space separation method for rejecting nearby interference in Meg Measurements. Physics in Medicine and Biology, 51(7), 1759–1768. 10.1088/0031-9155/51/7/008

86. Ulanovsky, N., Las, L., & Nelken, I. (2003). Processing of low-probability sounds by cortical neurons. Nature Neuroscience, 6(4), 391–398. 10.1038/nn1032

87. Vecchi, T., & Gatti, D. (2020). Memory as prediction: From looking back to looking forward. The MIT Press.

88. Wacongne, C., Changeux, J.-P., & Dehaene, S. (2012). A neuronal model of predictive coding accounting for the mismatch negativity. The Journal of Neuroscience, 32(11), 3665–3678. 10.1523/jneurosci.5003-11.2012

89. Woolrich, M. W., Hunt, L., Groves, A., & Barnes, G. (2011). Meg beamforming using Bayesian PCA for adaptive data covariance matrix regularization. NeuroImage, 57(4), 1466–1479. 10.1016/j.neuroimage.2011.04.041

90. Woolrich, M. W., Jbabdi, S., Patenaude, B., Chappell, M., Makni, S., Behrens, T., Beckmann, C., Jenkinson, M., & Smith, S. M. (2009). Bayesian analysis of neuroimaging data in FSL. NeuroImage, 45(1). 10.1016/j.neuroimage.2008.10.055

